# Overlapping activities of ELAV/Hu RNA binding proteins specify multiple neural alternative splicing programs

**DOI:** 10.1101/2020.09.21.305912

**Authors:** Seungjae Lee, Binglong Zhang, Lu Wei, Raeann Goering, Sonali Majumdar, J. Matthew Taliaferro, Eric C. Lai

**Affiliations:** Department of Developmental Biology, Sloan Kettering Institute, New York, NY 10065, USA; Department of Biochemistry and Molecular Genetics, University of Colorado Anschutz Medical Campus, Aurora, CO 80045, USA; RNA Bioscience Initiative, University of Colorado Anschutz Medical Campus, Aurora, CO 80045, USA

## Abstract

ELAV/Hu factors are conserved RNA binding proteins that play diverse roles in mRNA processing and regulation. The founding member, *Drosophila* Elav, was recognized as a vital neural factor 35 years ago. Nevertheless, still little is known about its impacts on the transcriptome, and potential functional overlap with its paralogs. Building on our recent findings that neural-specific lengthened 3’ UTR isoforms are co-determined by ELAV/Hu factors, we address their impacts on splicing. In ectopic contexts, all three members (Elav, Fne and Rbp9) induce similar and broad changes to cassette exon and alternative last exon (ALE) splicing. Reciprocally, double mutants of *elav/fne*, but not *elav* alone, have opposite effects on both types of mRNA processing events in the larval CNS. Accordingly, while *fne* mutants are normal, *fne* loss strongly enhances *elav* mutants with respect to neuronal differentiation. While manipulation of *Drosophila* ELAV/Hu factors induces both exon skipping and inclusion, motif analysis indicates their major direct effects are to suppress cassette exon usage. Moreover, we find direct analogies in their roles in global promotion of distal ALE splicing and terminal 3’ UTR extension, since both involve local suppression of proximal polyadenylation signals via ELAV/Hu binding sites downstream of cleavage sites. Finally, we provide evidence for analogous co-implementation of distal ALE and APA lengthening programs in mammalian neurons, linked to ELAV/Hu motifs downstream of regulated polyadenylation sites. Overall, ELAV/Hu proteins orchestrate multiple conserved programs of neuronal mRNA processing by suppressing alternative exons and polyadenylation sites.

## Introduction

The vast majority of genes in higher eukaryotes are subject to a variety of alternative processing mechanisms that diversify the functional outputs of the transcriptome [1,2]. The usage of alternative promoters, engagement of distinct internal and/or last exons by alternative splicing, and the utilization of alternative polyadenylation signals (PAS), can collectively generate transcript isoforms that differ in 5’ UTRs, coding exons, and/or 3’ UTRs (**Figure 1**). These regulatory regimes have broad consequences for differential regulation of isoforms as well as to broaden the protein outputs of an individual locus, and are aberrant in disease and cancer [3,4].

**Figure 1.**
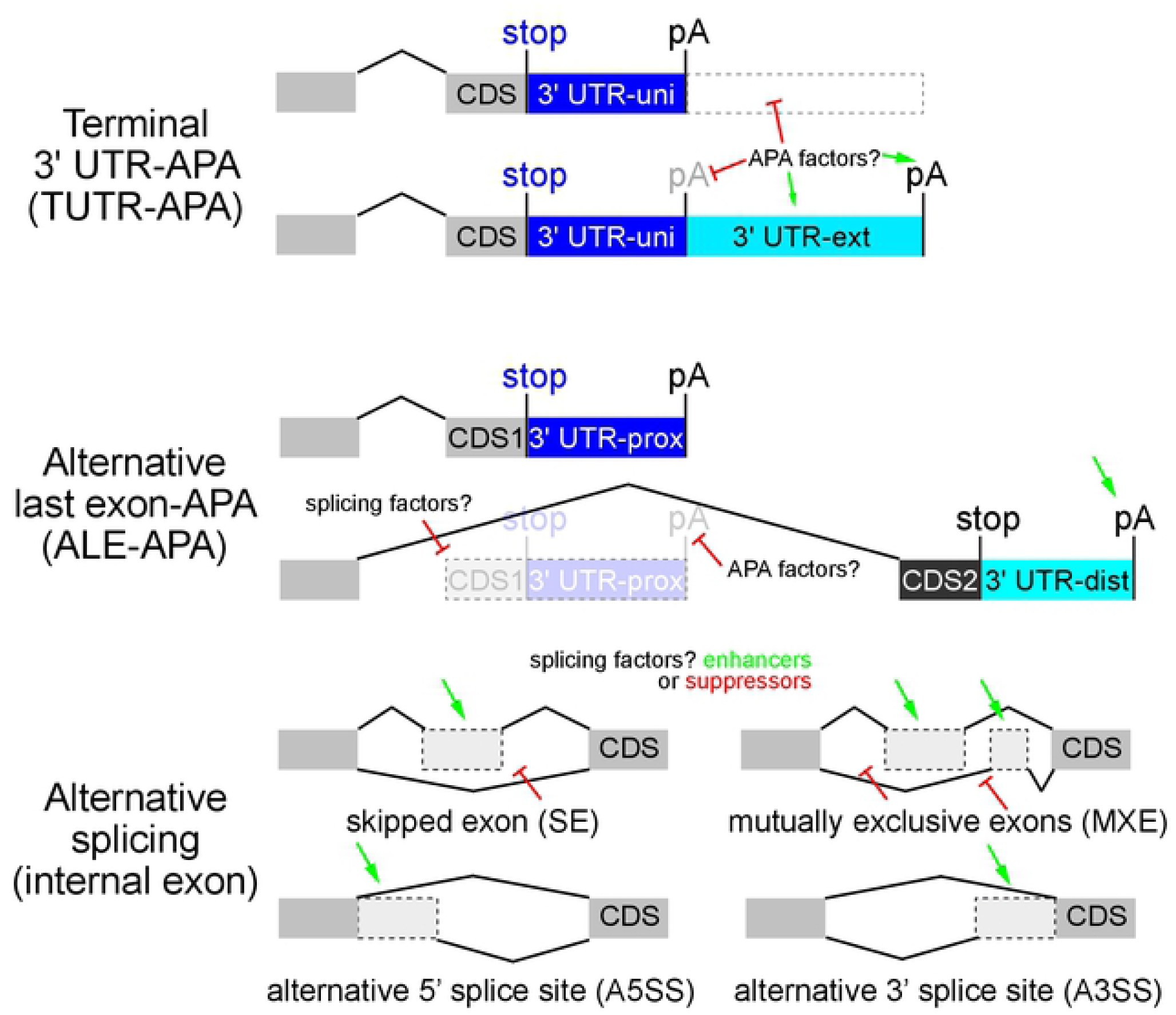
Summary of alternative polyadenylation and alternative splicing. Top, Alternative polyadenylation (APA) within the terminal 3’ UTR (TUTR-APA) generates 3’ UTR isoforms of variable lengths (universal/uni vs. extension/ext). Amongst other mechanisms, APA factors might modulate recognition of proximal pA signals, or selectively modulate the stability of ext isoforms. Middle, Alternative last exon (ALE)-APA generates isoforms with variable coding last exons, along with entirely distinct 3’ UTRs (proximal/prox vs. distal/dist). In principle, these could be regulated by splicing recognition, or by pA site recognition. Bottom, Alternative splicing of internal exons can be classified into several groups, and can be influenced by splicing enhancer or suppressors that bind in exonic or intronic regions. Not shown, alternative 5’ exons generated by distinct transcriptional start sites. This study will focus on exon skipping and the mechanistic relationship of TUTR-APA and ALE-APA.

While all of these regulatory concepts are applicable to all tissues and cells, the nervous system is well-known to exhibit particularly exuberant deployment of alternative isoforms [5,6]. For example, vertebrate brains encode the most diverse and conserved sets of alternative splicing events [7,8]. An extreme example is alternative splicing of the *Drosophila* immunoglobulin superfamily gene *Dscam*, which can generate ∼38,000 distinct proteins [9]. Indeed, experimental manipulations demonstrate that a high diversity of isoforms is functionally required for neuronal wiring [10]. The nervous system also exhibits broad usage of other unusual splicing programs. For example, mammalian brains preferentially express a suite of short “microexons” (encoding <10aa), which appear to be dysregulated in neurological disease [11]. Back-splicing that generates circular RNAs can be considered as another type of alternative splicing, and this class of species is most abundant and diverse in the nervous system [12,13]. Finally, *Drosophila* and vertebrate CNS exhibit the longest 3’ UTRs of any tissue [14-16].

Splicing and polyadenylation are catalyzed by large multisubunit RNA processing complexes, the former by the spliceosome and the latter by the cleavage and polyadenylation (CPA) machinery. As may be expected, direct experimental manipulations of core spliceosome or polyadenylation factors have broad impacts to deregulate isoform choice. The levels of many core splicing and polyadenylation factors differ across development and/or tissue or cell type, suggesting that their endogenous modulation may in part be linked to alternative mRNA processing. Reciprocally, trans-acting factors can impinge on the activities of splicing or polyadenylation machineries, either positively or negatively, to confer alternative processing.

The diversity of molecular processing mechanisms is best understood for alternative splicing [1,17]. Most of these strategies influence the definition of exons or introns, resulting in differential inclusion or exclusion of sequences in mature mRNAs (**Figure 1**). The SR family of RNA binding proteins (RBPs) provide a paradigm for proteins that bind exonic sequences to promote their inclusion, but certain SR factors bind intronic sequences to promote exon exclusion [18]. Many other non-SR proteins can regulate splicing (e.g. Nova, ELAV/Hu factors, PTBP proteins, MBNL proteins, etc.), and they generally mediate their effects by binding in the vicinity of splice sites. Notably, several of these have position-specific effects. For example, RBP binding overlapping a splice site can block exon inclusion, but intronic binding of the same factor can also promote exon inclusion [19-22].

Regarding APA, there are numerous ways that core CPA and trans-acting factors that influence sites of 3’ cleavage, and these can have distinct impacts depending on location within the gene [2,4]. When multiple pA sites are differentially utilized in the same terminal 3’ UTR exon (“TUTR-APA”, also referred to as tandem APA), this results in inclusion or exclusion of cis-regulatory sites within nested 3’ UTRs (**Figure 1**). The length of alternative 3’ UTRs can be quite substantial (>10 kb), and there may be sequence-independent effects of 3’ UTR length variants. Another general class of APA events occur within regions that are internal to the last exon of the longest gene model. These might occur within internal exons or introns, and have variously been termed upstream region-APA, intronic APA, or alternative last exon (ALE)-APA.

If internal polyadenylation impairs the stability of the alternative transcript, this can result in loss-of-function of that isoform. However, ALE-APA can also produce alternative stable transcripts, which encode distinct protein isoforms (**Figure 1**). In some cases, internally polyadenylated isoforms yield proteins that lack C-terminal domains of “full-length” counterparts, which might inactivate them and/or encode dominant negative or neomorphic activities [23]. On the other hand, there are many examples of loci with distinct ALE isoforms that harbor different activities, analogous to distinct internal splice isoforms. Indeed, early characterizations of alternative polyadenylation involved ALE isoforms that switch the localization of IgM heavy chain products from secreted to membrane-bound [24], or generate distinct protein isoforms of calcitonin/CGRP[25]. Beyond distinct coding potential, different ALE isoforms can also be subject to regulated subcellular localization, particularly within neurons [26-28]. Although less is understood about trans-acting factors that influence APA, several RBPs (e.g. PABPN1, CELF1, etc.) operate in a conceptually similar manner to splicing regulators. Namely, several RBPs modulate 3’-end selection by binding in the vicinity of cleavage sites to oppose the action of the CPA machinery [29-33].

The ELAV/Hu family of RBPs are conserved across metazoans and play broad roles in RNA biogenesis, including alternative splicing, APA, target stability, translation, and localization [34-41]. In *Drosophila*, Elav was proposed as a direct regulator of both splicing and APA. Notably, there is a conceptual mechanistic link between these regulatory processes. The first direct target of Elav characterized is *erect wing* (*ewg*), and it controls splicing of its alternative last exons [40]. Elav also regulates the splicing of certain internal exons [42,43], but unlike other strategies for splicing control mentioned above, Elav controls neural-specific *ewg* by modulating cleavage and polyadenylation. In particular, Elav suppresses cleavage at the 3’ end of an internal *ewg* terminal exon isoform, thereby permitting transcription to extend to the distal terminal exon. This mechanism also applies to neural-specific of *neuroglian* (*nrg*) [41]. An analogous function for Elav was reported to promote neural-specific 3’ UTR lengthening at select genes bearing tandem polyA signals within the same 3’ UTR [38]. In this setting, it was proposed that association of Elav with a proximal polyA signal inhibits processing by the CPA machinery, thereby permitting transcription into distal 3’ UTR segments.

These studies focused on regulation of a few specific *Drosophila* genes, but broader impacts of Elav and/or its paralogs on the mRNA processing were not yet addressed. In work submitted, we made the following observations [44]. First, although Elav has long been considered to be embryonic lethal, we found that *elav* deletion mutants are viable as 1st instar larvae. With access to elav null larval CNS we found that its complete loss was compatible with expression of many neural APA 3’ UTR extensions. Second, we found using gain-of-function strategies, that all three *Drosophila* ELAV/Hu members (Elav, Fne, Rbp9) have similar capacities to induce a neural 3’ UTR extension landscape in an ectopic setting (S2 cells). Third, we found that functional redundancy is endogenously relevant, because elav/fne double mutant larval CNS exhibit a severe loss of neural 3’ UTR extension landscape. Fourth, the functional overlap of Elav and Fne involves a regulatory interplay, because Elav represses *fne* alternative splicing that switches it from a cytoplasmic to a nuclear isoform.

In this study, we take advantage of these gain-of-function and loss-of-function genomic datasets to study the impact of *Drosophila* ELAV/Hu factors on alternative splicing, including both internal exons as well as terminal exons. We broaden the set of cassette exons and alternative 5’ or 3’ splice sites that are regulated by Elav and Fne. Moreover, we define a broad program of neural ALE splicing that is specified by Elav or by overlapping activities of ELAV/Hu factors. Genomic analyses reveal mechanistic parallels between neural ALE splicing and neural 3’ UTR lengthening, demonstrating that these are analogous processes that operate in a directional manner on transcripts to promote the inclusion of distal exonic sequences in neurons. Consistent with the unexpected functional overlap of Elav and Fne, we provide evidence for strong phenotypic enhancement of *elav* mutants by *fne* mutation in the context of larval neuronal differentiation. Finally, we extend these findings to mammals, and provide evidence for coincident shifts towards usage of distal ALE isoforms and extended tandem 3’ UTRs in mammalian brains and during directed neuronal differentiation. These regulatory programs exhibit similar signatures for direct regulation by ELAV/Hu proteins at bypassed pA sites, suggesting conservation and coordination of these two RNA processing pathways across metazoan neurons.

## Results

### Broad impacts of *Drosophila* ELAV/Hu factors on alternative splicing

Mammalian ELAV/Hu factors are documented regulators of alternative splicing, mediating both exon inclusion and exclusion. This applies to both ubiquitously expressed HuR [45,46] as well as neuronally restricted HuB/C/D [19,47]. However, even though *Drosophila* Elav was shown to be involved in splicing [48] before its mammalian counterparts [49], the internal splicing of only two *Drosophila* genes (*Dscam1* and *arm*) has been reported to be dependent on Elav [42,43]. Moreover, the capacity of other *Drosophila* ELAV/Hu paralogs (Fne and Rbp9) to influence splicing is largely unknown, but presumed not endogenously relevant due to their cytoplasmic localization [50,51].

We investigated if these RBPs have broader impacts on alternative splicing. We first generated RNA-seq datasets from S2 cells that ectopically expressed wt Elav/Fne/Rbp9, or RNA binding-defective variants bearing inactivating point mutations in all three RRM domains (also referred to as “mut”). We then used rMATS[52] to investigate various classes of alternative splicing; this package classifies alternative internal isoforms (**Figure 1**). This revealed 610 differentially spliced exons across all alternative splice types, and relatively comparable numbers of exons exhibited gains or losses of usage in the presence of ectopic ELAV/Hu factors (**Supplementary Figure 1**). Exon skipping comprised the major deregulated category, and these exhibited mild directional bias for exclusion by ectopic wt ELAV/Hu factors compared to their mutant variants, but substantial numbers of exons were driven towards inclusion (**Supplementary Figure 1**). Since cassette exons were the dominant source of alternative splicing, we performed principal components analysis (PCA) on exon skipping. This showed that mutant ELAV/Hu datasets clustered near control S2 cells, while wildtype Elav/Fne/Rbp9 were well-separated (**Figure 2A**). Therefore, ectopic ELAV/Hu proteins induced substantial splicing variation dependent on their RNA binding activities. We observed substantial overlap in the regulatory influences of these factors (examples of individual genes shown in **Supplementary Figure 2**). However, as we only had single RNA-seq datasets for each, we focused on cassette exons that exhibited similar differential splicing in the presence of all three wildtype ELAV/Hu RBPs in S2 cells (**Figure 2B-C**).

**Figure 2.**
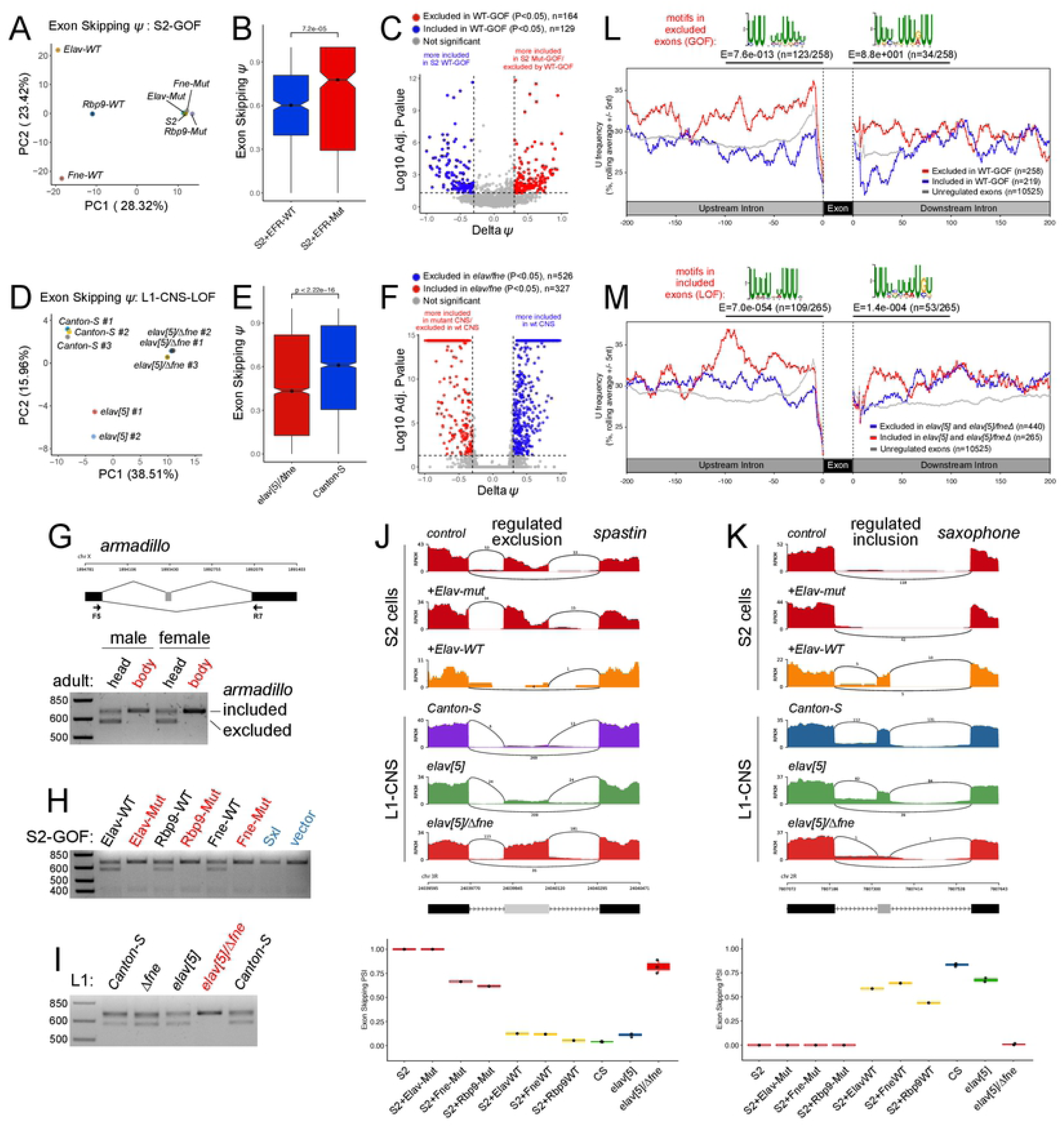
Overlapping activities of *Drosophila* ELAV/Hu RBPs control alternative cassette exon splicing. (A) Principal components analysis (PCA) of alternative splicing of cassette exons (percent spliced in, PSI/*Ψ* values) RNA-seq data from S2 cells transfected with wildtype (wt) or 3xRRM-mut (mut) versions of Elav, Fne and Rbp9. (B) Fraction of exon skipping events in combined S2-Elav/Fne/Rbp (EFR)-wt or -mut transfection datasets. (C) Volcano plot of alternative spliced cassette exon events in S2 cells following manipulation of ELAV/Hu RBPs. Note that exons that are preferentially excluded by wt-ELAV/Hu RBPs correspond to exons that are preferentially included in control S2 cells, or cells expressing mutant ELAV/Hu RBPs, and *vice versa*. (D) PCA of cassette exon splicing in L1-CNS RNA-seq data from *Canton-S, elav* mutant, and double *elav/fne* mutants. (E) Fraction of exon skipping events in *elav/fne* mutant and *Canton-S* L1-CNS. (F) Volcano plot of alternatively spliced cassette exon events in *Canton-S* and *elav/fne* mutant L1-CNS. (G-I) rt-PCR validation of an alternatively spliced cassette exon in *armadillo*. (G) The exon-excluded isoform is preferentially expressed in the nervous system, and is thus detected in adult heads but not bodies. (H) In S2 cells, the exon-excluded isoform of *armadillo* is induced by Elav/Rbp9/Fne-wt, but not their mutant counterparts or the similar RBP Sxl. (I) The exon-excluded *armadillo* isoform is present in L1-CNS of wildtype (*Canton-S*) and single mutants of *fne* and *elav*, but not the double mutant. (J) Example of a novel exon exclusion target of ELAV/Hu RBPs. (K) Example of a novel exon inclusion target of ELAV/Hu RBPs. (L) Global analysis of exons excluded by misexpression of Elav/Fne/Rbp (EFR) in S2 cells shows U-enrichment in flanking intronic regions, especially in the upstream regions. This is not seen in flanking introns of exons included by EFR. De novo motif analysis shows the top enrichment sequence matches the ELAV/Hu binding site. (M) Global analysis of exons included in *elav* and *elav/fne* mutant L1-CNS, compared to *Canton-S*, shows similar positional enrichment of intronic uridine and Elav binding sites.

Since the S2 cell system is an ectopic setting, we addressed the endogenous impact of ELAV/Hu factors on splicing using recently available RNA-seq data from L1-CNS [44]. We found that the first larval instar (L1)-CNS is a setting in which there is substantial functional overlap of Elav and Fne to direct neural 3’ UTR extensions, especially as a consequence of nuclear re-localization of Fne in *elav* mutants [44]. Interestingly, this involves a regulatory interplay of these factors, since Elav represses a novel alternative splice isoform of *fne* that is nuclearly localized. As Fne is neural-specific, this regulatory paradigm could not be appreciated from S2 cells. Since we were particularly interested in identifying endogenous splicing targets confidently, which may include other neural-restricted loci, we utilized replicate datasets for these *in vivo* samples. Accordingly, we examined global alternative splicing in L1-CNS from control *Canton-S* (wildtype), *elav[5]* null, and *elav[5]/fne[Δ]* double deletion backgrounds.

As with the S2 analyses, categorization of all alternative splicing events showed that exon skipping compromised the major class (**Supplementary Figure 1**). In addition, while *elav* deletion clearly had impact on neural splicing, all effects were more profound in *elav/fne* double mutants. This could be visualized by the greater numbers of affected loci when comparing either *elav* or *elav/fne* to control, or by comparing the single and double mutants (**Supplementary Figure 1**). Of note, the breadth of splicing deregulation in the nervous system was even greater than in the ectopic studies. We detected 1624 splicing alterations in double mutants, with cassette exons again representing the majority of events (**Supplementary Figure 1**). PCA of alternative cassette exon usage across L1-CNS replicates highlights the more severe effects of dual loss of *elav/fne* compared to *elav* (**Figure 2D**). We illustrate the impacts of Elav/Fne double mutants on exon skipping in **Figure 2E-F**, and focus on these loci for further analyses. Overall, both ELAV/Hu factors are required to maintain endogenous neural splicing.

We selected *armadillo* (*arm*) for further evaluation, as it was previously known to yield a neural-specific splice isoform [53] and comprises one of two known cassette exon targets of Elav [42]. However, we note that previous studies did not demonstrate endogenous regulation of *arm* splicing *in trans*. The shorter neural-specific isoform can be easily detected by rt-PCR in adult heads, but not in bodies (**Figure 2G**). Not only is this *arm* exon excluded by ectopic Elav in S2 cells, it is similarly excluded by ectopic Fne and RBP9, but not by any of their 3xRRM-mut counterparts (**Figure 2H**). In contrast, overexpression of the closest relative of ELAV/Hu members in *Drosophila*, Sex-lethal (Sxl), did not induce neural *arm* splicing, even though ectopic Sxl can regulate splicing of an Elav target reporter [54]. This provides evidence for the specificity of ELAV/Hu factor-mediated mRNA processing. We next analyzed L1 samples and observed that accumulation of the neural *arm* isoform was relatively unaffected in either *elav* or *fne* single deletion mutants. However, rt-PCR from *elav/fne* double mutant larvae showed complete absence of the neural *arm* isoform (**Figure 2I**). Therefore, both Elav and Fne are needed to generate neural *arm* splicing.

We highlight other examples of splicing dysregulation following manipulation of ELAV/Hu factors. More than 250 alternatively spliced cassette exons were collectively excluded in S2-GOF experiments, and reciprocally, a similar number became ectopically included in L1-CNS-LOF mutants. Examples of genes with exon-skipping promoted by ELAV/Hu factors include the Dpp pathway transcription factor *Medea* and the microtubule severing factor *spastin*. Like *arm*, these loci harbor exons that are included in control S2 cells or cells expressing 3xRRM-mutant versions of Elav/Fne/Rbp9, but are efficiently skipped in cells expressing wildtype Elav/Fne/Rbp9 (**Figure 2J** and **Supplementary Figure 2**). On the other hand, their exons are skipped in wt and *elav[5]* L1-CNS, but are included in *elav[5]/fne[Δ]* L1-CNS (**Supplementary Figure 2**). We also observe genes with the reciprocal behavior. For instance, the BMP receptor *saxophone* (**Figure 2K**), and two genes involved in cytoskeleton remodeling, *LIM kinase 1* and the cysteine peptidase inhibitor *sigmar* (**Supplementary Figure 2**), exemplify loci with cassette exons whose inclusion is promoted by ELAV/Hu factors, and have reciprocal behavior in GOF and LOF settings. The numbers of these genes with this splicing response are even greater (**Supplementary Figure 1**).

Taken together, these analyses reveal that beyond a couple of loci studied over the past 20 years [42,43], *Drosophila* ELAV/Hu factors are global regulators of alternative splicing of cassette exons in neurons. This realization depends on analysis of both sufficiency data as well as the appropriate double mutant data.

### Exons excluded by *Drosophila* ELAV/Hu factors exhibit signatures of direct regulation

Prior analyses in mammals detected numerous alternative splicing events regulated by HuR [45,46] or neuronal Hu proteins [19], but were on the whole equivocal as to the layout of ELAV/Hu-regulated exons. Based on independent analyses of a few dozen alternative splicing events, it was concluded that binding of ELAV/Hu RBPs is enriched upstream of both included and excluded exons [19,46], but it was also suggested that many alternative splicing events detected after HuR depletion were likely indirect effects [46].

We addressed this in our data by analyzing sequence properties and performing *de novo* motif searches amongst different cohorts of alternatively spliced cassette exons. We observed overtly distinct nucleotide distributions depending on whether they were excluded or included, and these exhibited reciprocal features in GOF and LOF data. In particular, exons that were preferentially excluded in S2-ELAV/Hu-GOF data exhibited distinctly elevated U-content in flanking upstream intronic regions, while this was also the case for exons that became preferentially included in *elav/fne* mutant L1-CNS (**Figure 2L-M** and **Supplementary Figure 3**). Importantly, even though these regions are expected to have biased content because they overlap polypyrimidine (C/U) tracts located upstream of strong splice acceptor sequences, comparable elevation of U-content was not observed upstream of exons that displayed opposite regulatory behavior in S2 or L1-CNS. Instead, we observed elevated A-content downstream of exons that were preferentially included in S2-ELAV/Hu-GOF settings, or were excluded in *elav/fne* mutant L1-CNS (**Supplementary Figure 3**). Therefore, elevated U-content in flanking introns is a specific feature of exons that are excluded by ELAV/Hu factors.

**Figure 3.**
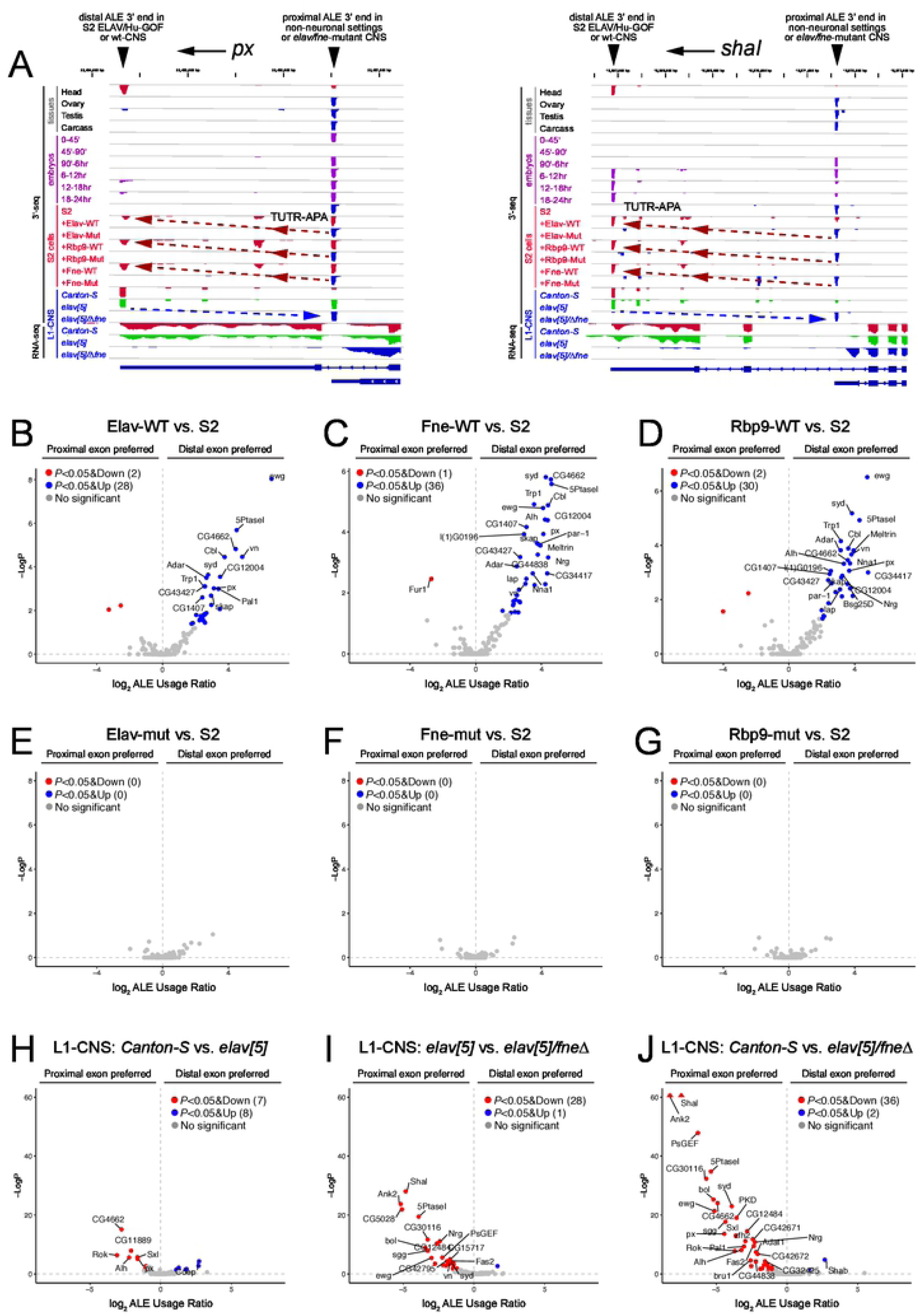
Overlapping activities of *Drosophila* ELAV/Hu RBPs control alternative last exon (ALE) splicing. (A) Examples of neural-specific distal ALE splice isoforms whose utilization exhibits necessity and sufficiency for ELAV/Hu RBPs. *px* and *ShaI* exemplify distal ALE isoforms that are (1) preferentially expressed in head compared to other tissues, (2) are developmentally induced during the timecourse of embryogenesis, (3) are induced in S2 cells upon transfection of wildtype Elav/Rbp9/Fne but not their RRM-mutant (Mut) counterparts, and (4) are expressed in dissected wildtype *Canton-S* and *elav[5]* null L1-CNS, but not in *elav[5]/Δfne* double mutant L1-CNS. Tracks are labeled as to 3’-seq or RNA-seq data. (B-D) Global analysis of misexpression of wildtype ELAV/Hu RBPs in S2 cells shows highly directional induction of distal ALE isoforms. (E-G) Misexpression of RRM-mutant ELAV/Hu RBPs in S2 cells does not alter ALE isoform usage. (H-I) Combined deletion of *elav* and *fne* causes a global reversion of distal-to-proximal ALE isoform usage switch in L1-CNS.

We performed motif discovery on these cohorts of introns, noting that the conserved binding site of ELAV/Hu factors is known to be U-rich [55]. Strikingly, close matches to the empirically-determined binding sites for Elav/Fne/Rbp9 [55] are highly enriched upstream of exons that were ectopically excluded in S2-GOF data, and reciprocally in exons that were ectopically included in elav/fne mutant L1-CNS data (**Figure 2L-M**). These were the most frequent (∼40-50% of regulated exons) and most significant motifs found in the intronic regions upstream of exons regulated in these directions; they were also significantly enriched downstream of these regulated exons, albeit less frequently (**Figure 2L-M**). By contrast, ELAV/Hu-type binding sites were not enriched in the vicinity of included exons. A comprehensive summary of *de novo* motif searches are presented in **Supplementary Figure 3**, and include an A-rich motif downstream of exons that are preferentially included in the presence of ELAV/Hu proteins in either S2 (WT vs. Mut overexpression) or L1-CNS (WT vs. *elav/fne* double mutant) settings.

In summary, these data indicate that *Drosophila* ELAV/Hu proteins mediate an extensive set of direct and indirect effects on alternative splicing. Our analyses depend on sufficiency tests that demonstrate similar capacities of multiple ELAV/Hu members, which led us to recognize endogenous functional overlap by Elav and Fne to specify the neural transcriptome. The major direct consequences of *Drosophila* ELAV/Hu RBPs comprise exclusion of cassette exons, via binding sites in flanking intronic regions that are preferentially located upstream of regulated exons. We also observe that *Drosophila* ELAV/Hu RBPs are necessary and sufficient to induce a second layer of likely indirect regulatory consequences, which are enriched for exon inclusion events and A-rich motifs located downstream of regulated exons. It remains to be determined what might regulate this program, but we note that PABP, PABP2 (PABPN1), ZC3H14/dNab2, and hnRNP-Q (Syncrip) proteins associate with qualitatively similar A-rich motifs [55-57].

### Overlapping activities of ELAV/Hu factors also drive global distal ALE splicing

A special class of regulated splicing occurs at alternative last exons (ALEs, **Figure 1**). As noted, two genes (*ewg* and *nrg*) were known to have Elav-dependent neural ALE splicing [41,48], and ectopic Elav/Fne/Rbp9 have common abilities to promote distal ALE switching of *nrg* [54]. To gain broader insight into regulation of ALE by ELAV/Hu factors, we exploited 3’-seq datasets from S2-GOF and L1-CNS-LOF conditions [44], which permit more precise quantification of 3’-isoform shifts than RNA-seq data [58].

We illustrate newly recognized examples of these loci in **Figure 3A** and **Supplementary Figure 4**. In particular, *px* and *shaI* embody relevant expression and regulatory principles (**Figure 3A**). First, when sampling wildtype tissue data (head, ovary, testis, and carcass) or across a series of 6 timepoints that span embryonic development, we see that these genes preferentially or selectively express the distal ALE isoforms in the head or in late embryo stages when the nervous system has begun to differentiate. Second, comparing S2 cells that express wildtype or RRM-mutant ELAV/Hu factors, we see that all wildtype conditions and no mutant conditions are associated with ectopic induction of distal ALE isoforms. Third, we see that the distal ALE isoforms are present in wildtype and *elav* null L1-CNS, but revert to proximal ALE isoforms in *elav/fne* double mutant L1-CNS (**Figure 3A**).

**Figure 4.**
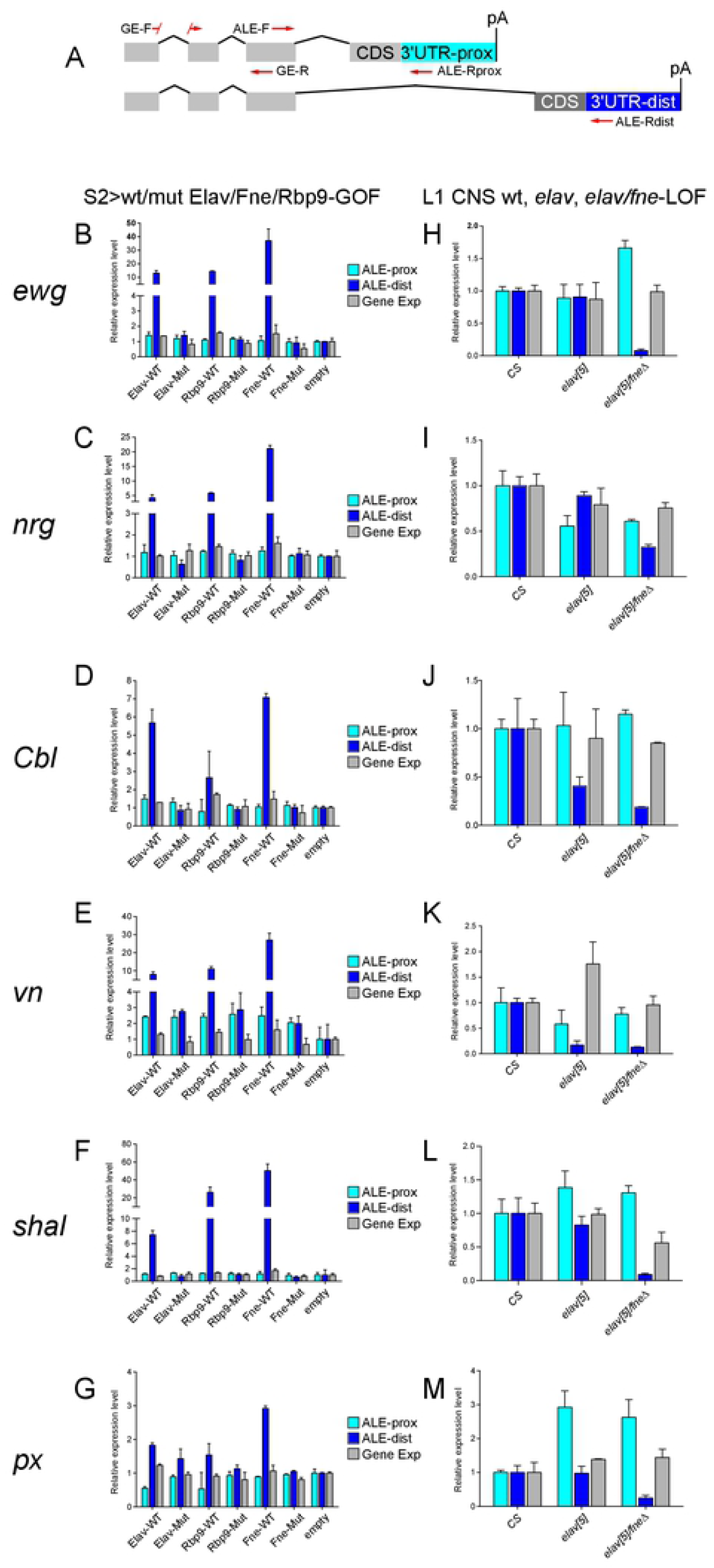
Experimental validation of distal ALE switching mediated by *Drosophila* ELAV/Hu RBPs. (A) Schematic of ALE isoforms and qPCR primers to assess gene expression (GE), proximal ALE isoform, and distal ALE isoform expression. Data were normalized to *rpl32*. (B-G) Assessment of 6 ALE targets in S2 cells upon misexpression of wildtype (WT) or 3xRRM-mutant (Mut) forms of Elav, Rbp9 and Fne. All ELAV/Hu RBPs induce distal ALE isoforms, dependent on their RNA binding capacity, with relatively little effects on total locus expression. (H-M) Assessment of the same ALE targets in L1-CNS upon endogenous deletion of *elav* or both *elav/fne*. Certain targets exhibit sensitive distal ALE loss in single mutant conditions (e.g. *Cbl* and *vn*), but all of these loci exhibit decreased distal ALE isoform expression in the double mutant.

Since rMATS does not evaluate alternative last exon (ALE) splicing, we utilized our 3’-seq analysis pipelines [59] to assess ALE switching. Notably, ectopic Elav/Fne/Rbp9 all induced global shifts in ALE splicing in S2 cells (**Figure 3B-D**). These changes were quantitatively robust, affected a largely overlapping set of genes, and caused a directional shift towards distal 3’ UTR usage. In addition, these effects were fully dependent on their RNA-binding capacities, since their all 3xRRM-mutant versions were relatively inert (**Figure 3E-G)**.

Based on this, we assessed the endogenous requirement of ELAV/Hu factors for distal ALE switching. Notably, Elav is required for neural distal ALE switching of *ewg* and *nrg* in photoreceptors, while *fne/rbp9* were reported to be dispensable for this process, either in photoreceptors or more generally in third instar larval or adult CNS [54]. However, ALE deployment has not been examined in first instar larval CNS. Perhaps surprisingly, we observed almost no ALE changes in these genes between control and *elav[5]* mutants (**Figure 3H**), even though the neurons in these samples have definitively been null for Elav for 12-18 hours.

However, we observed substantial shifts towards proximal ALE isoform usage in *elav[5]/Δfne* double mutants; the differences were greater than when comparing *Canton-S* to *elav[5]* (**Figure 3I-J**). Overall, we generally observed reciprocal behavior between the GOF and LOF datasets with respect to alteration of distal ALE isoform usage (**Figure 3** and **Supplementary Figure 4**), indicating that combined activity of ELAV/Hu RBPs plays a broad endogenous role to induce distal ALE isoforms in the larval CNS.

We validated these genomic data using quantitative rt-PCR for universal exon, proximal 3’ UTR and distal 3’ UTR amplicons of ALE loci (**Figure 4A**). With such assays, we could quantify changes in ALE isoforms as well as determine overall changes in gene expression. To validate our S2 samples, we selected additional neural-extension APA targets from our genomic data [44] and observed that tandem 3’ UTR extensions of *tai* and *ctp* were specifically induced upon expression of wildtype (but not 3xRRM-mutant) Elav/Fne/Rbp9 in S2 cells (**Supplementary Figure 5**). With these confirmations, we tested six ALE loci and observed that ectopic Elav/Rbp9/Fne all have RRM-dependent capacities to promote the accumulation of distal ALE isoforms (**Figure 4B-G**). In general, these isoform changes do not substantially affect total gene expression levels. We observed opposite effects in L1 larvae (**Figure 4H-M**). For the most part, single *elav[5]* null larvae did not exhibit ALE switching. This was particularly notable for *ewg* and *nrg*, which exhibit high dependency on Elav in the larval eye disc [41,48], but not in L1 larvae (**Figure 4H-I**). Other ALE loci such as *shal* and *px* were also not affected in *elav* mutant L1, but we did find two loci (*Cbl* and *vn*) with selective loss of distal ALE isoforms in this background (**Figure 4J-K**). However, the effects in *elav/fne* double mutant L1 larvae were much more severe. With the exception of *vn*, whose strong loss in the single mutant did not provide room for enhancement, *Cbl* exhibited further loss of distal ALE expression in the double while the other four genes exhibit synthetic phenotypes in the double mutant (**Figure 4H-M**).

**Figure 5.**
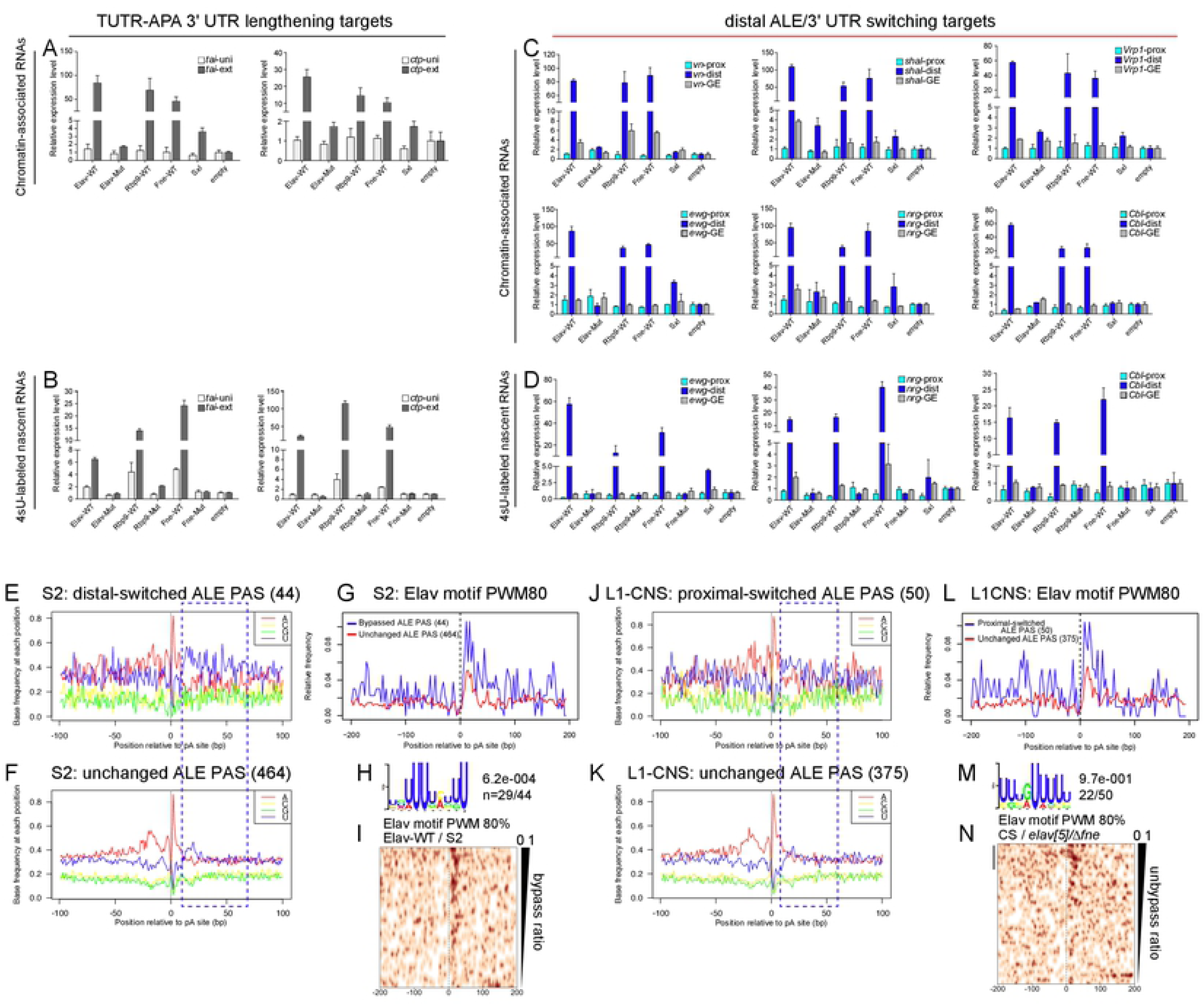
Mechanism of ALE-APA control by ELAV/Hu RBPs is shared with TUTR-APA targets. (A-B) Validation of new terminal 3’ UTR-APA (TUTR-APA) targets in the S2-GOF system. The 3’ UTR extensions of *tai* and *ctp* can be detected in chromatin-associated fractions (A) and in nascent transcripts isolated using 4sU labeling (B). (C) Distal ALE switching induced by GOF of ELAV/Hu RBPs also occurs in chromatin-associated fractions, but is not detected in RNA binding-defective Elav or with the homolog RBP Sxl. (D) Distal ALE switching induced by GOF of ELAV/Hu RBPs is detected in newly-synthesized transcripts. (E-I) Nucleotide and motif analysis of regions surrounding proximal ALE polyadenylation (pA) sites in S2 cells. (E-F) Nucleotide frequency of ALE cleavage sites that are bypassed by ELAV/Hu RBP-GOF (E) or that are unchanged (F) shows enrichment of downstream uridine amongst regulated loci. (G) Elav binding sites (80% match to position weight matrix, PWM80) are enriched downstream of regulated proximal ALE pA sites. (H) De novo search shows the Elav binding site is the most frequent and most significant motif downstream of regulated ALE pA sites. (I) Elav binding sites are correlated with the frequency of bypass as measured by distal isoform induction in the presence of ectopic Elav in S2 cells. (J-N) Nucleotide and motif analysis of regions surrounding proximal ALE pA sites in L1-CNS. (J-K) Nucleotide frequency of ALE cleavage sites that are switched from distal to proximal usage by *elav/fne-*LOF (J) or that are unchanged (K) shows enrichment of downstream uridine amongst regulated loci. (L) Elav binding sites are enriched downstream of regulated proximal ALE pA sites. (M) De novo search shows the Elav binding site is the most frequent and most significant motif downstream of endogenously regulated ALE pA sites in L1-CNS. (N) Elav binding sites correlate with distal-to-proximal ALE isoform switching in *elav/fne* mutant L1-CNS.

Altogether, these GOF and LOF analyses broadly extend the role of ELAV/Hu factors in driving distal ALE splicing in *Drosophila* neurons, from just two targets to several dozen. Moreover, they affirm that Fne plays a critical role to maintain neural ALE splicing when Elav is absent.

### ELAV/Hu RBPs induce distal ALE usage by inhibiting pA site usage of internal isoforms

We investigated mechanisms for how ELAV/Hu family RBPs induce distal alternative last exon splicing (**Figure 1**). In principle, this could be regulated at the level of splicing into the proximal alternative exon (ALE), or by altering the stability of the different isoforms. However, in the case of *ewg* [40] and *nrg* [41], Elav binds downstream of cleavage sites for proximal ALE isoforms and inhibits their usage, thereby permitting further transcription and splicing into the downstream ALE isoform. Notably, we found broad evidence for an analogous model as to how ELAV/Hu proteins implement neural 3’ UTR lengthening [44]. Therefore, we applied our assays to test if these principles might apply more broadly to other ALE targets, and across other members of the ELAV/Hu RBP family.

We next used the controlled S2 cell system to assess whether ELAV/Hu-mediated distal ALE switching occurred in subcellular and/or temporal contexts that were consistent with the pA site bypass model. For this purpose, we compared the levels of proximal and distal ALE isoforms with common amplicons for a panel of loci, in response to all three wildtype ELAV/Hu RBPs, 3xRRM-mut variant(s), or Sxl as a control U-rich binding RBP. For comparison, we performed additional assays of newly-validated TUTR-APA 3’ UTR lengthening targets *tai* and *ctp* (**Supplementary Figure 5**) and confirmed that ELAV/Hu RBP-induced 3’ UTR extension occurs in chromatin-associated transcripts (**Figure 5A**) and in newly-synthesized RNAs isolated by 4sU labeling (**Figure 5B**). We obtained reliably similar and specific induction of a panel of distal ALE isoforms by wildtype ELAV/Hu RBPs (**Figure 5C-D**), indicating similar features of molecular regulation for 3’ UTR lengthening and distal ALE switching.

We then examined if there were characteristic motif features at the 3’ termini of genes subject to ELAV/Hu-induced distal ALE switching. We first examined PAS, and classified them as to whether they were at the termini of the internal or the distal ALE isoforms, or whether they were internal to 3’ UTRs (**Supplementary Figure 6A**). As a reference, we analyzed single-end S2 cell genes, which are expected to harbor strong PAS. We represent the frequency of consensus PAS motifs 10-30 nt upstream of the mapped cleavage sites from 3’-seq data in stacked bar plots. We compared this to PAS derived from two types of S2-expressed ALE gene models, those that collectively exhibited distal ALE usage upon gain-of-function of Elav/Fne/Rbp9 (38 genes, 44 3’ UTRs) and those that were insensitive to their action (367 genes, 464 3’ UTRs). We observed that the PAS quality of internal ALE 3’ termini was substantially lower than constitutive 3’ ends, implying that they are preferentially bypassable (**Supplementary Figure 6B**). However, there was no substantial difference between regulated and non-regulated ALE genes.

**Figure 6.**
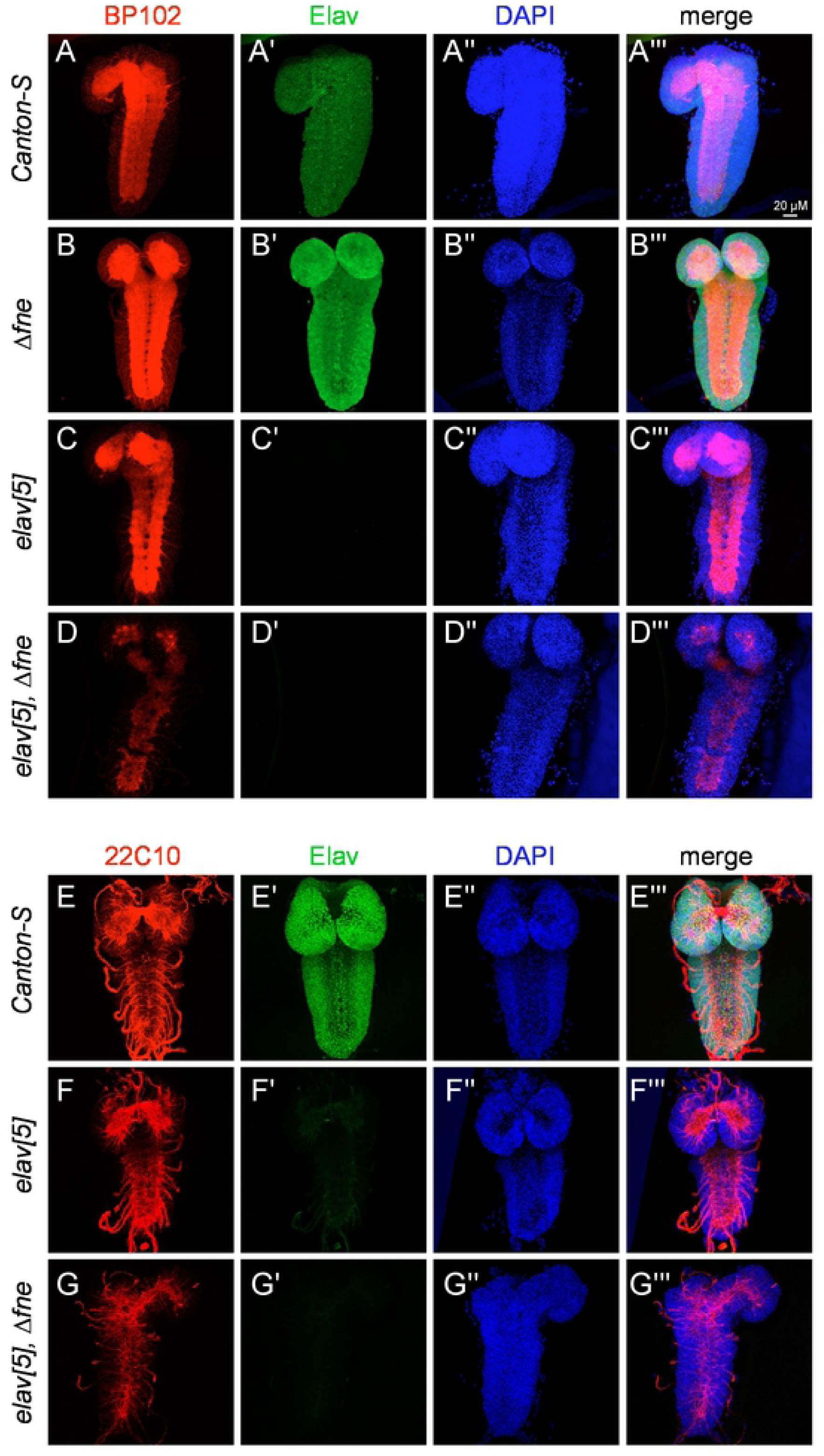
Combined activities of Elav and Fne are required for differentiation of the *Drosophila* CNS. Shown are stainings of the 1st instar larval central nervous system (L1-CNS). (A-D) Staining for BP102 (red), Elav (green) and DAPI (blue). (A-B) BP102 labels a ladder-like pattern of CNS commissures and connectives in control *Canton-S* (A) as well as *Δfne* (B). (C) *elav[5]* null mutant exhibits occasional breaks and midline crossing-over in BP102 labeling. (D) Double mutant *elav[5]/Δfne* exhibits strong reduction in BP102 signal and highly aberrant patterning including massive crossing-over and breaks. (E-G) 22C10 (Futsch) labels all axonal projections. (E) The characteristic pattern of normal 22C10 is shown in *Canton-S*. (F) *elav[5]* exhibit mild reduction and mispatterning of 22C10. (G) *elav[5]/Δfne* shows strong reduction in 22C10 signals and highly aberrant patterning.

We next examined for sequence characteristics in the vicinity of cleavage sites. We plotted the nucleotide distribution +/-100 nt of cleavage sites mapped by 3’-seq data, for internal ALE termini that were or were not sensitive to ELAV/Hu factors in S2 cells. Although the pattern was more noisy for the smaller ELAV/Hu-sensitive dataset, these plots indicated enhanced frequency of uridine downstream of their cleavage sites, compared to the control set (**Figure 5E-F**). This disparity was emphasized by plotting the uridine frequency downstream of cleavage sites between these sets of internal ALE termini (**Supplementary Figure 7**). Thus, although a U/GU-rich sequence downstream of pA sites generally contributes to cleavage by recruiting CstF, internal ALE termini that can be bypassed in the presence of ELAV/Hu factors exhibit distinct U-rich downstream context.

**Figure 7.**
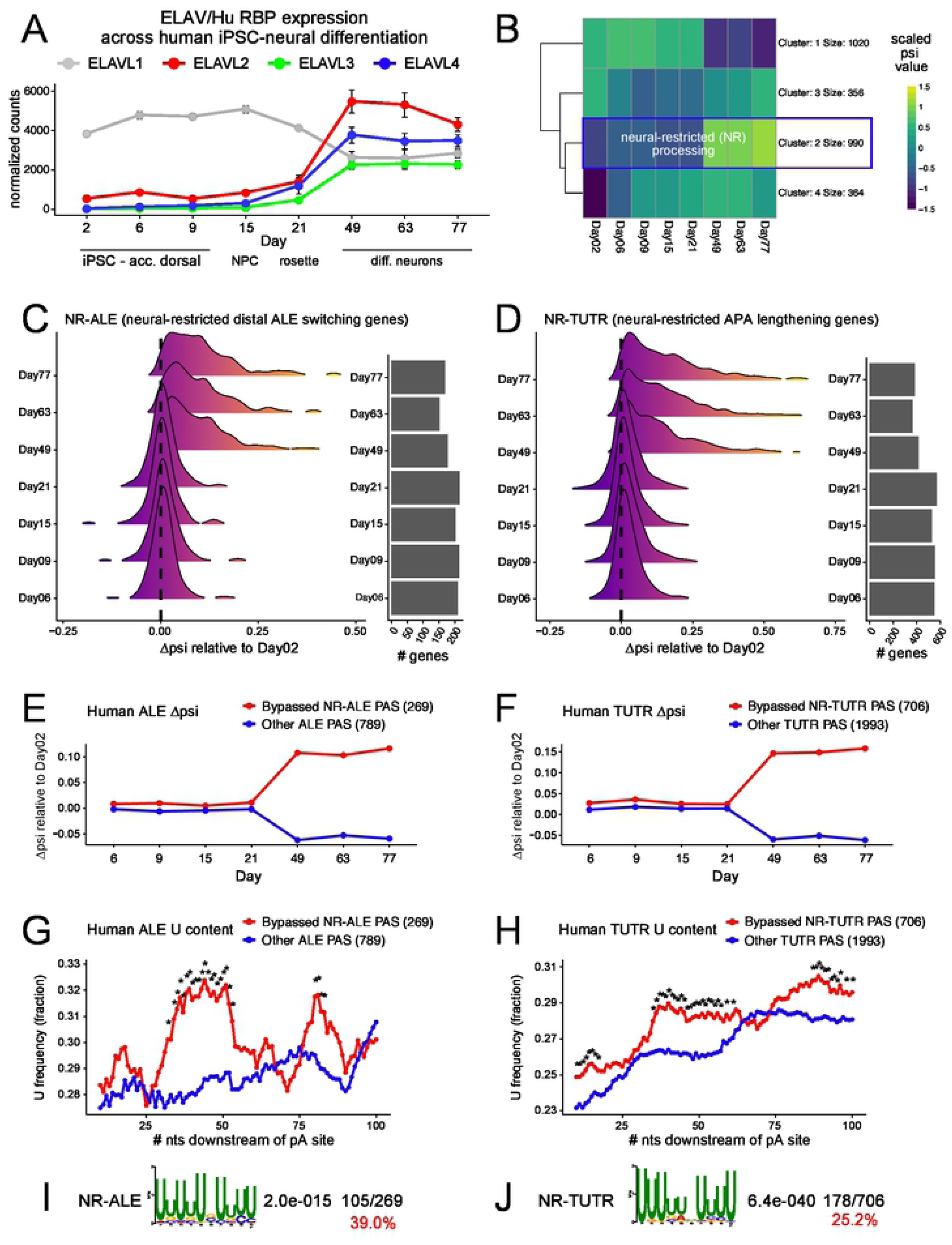
Evidence that mammalian neural ELAV/Hu RBPs are involved in distal ALE switching and 3’ UTR lengthening in neurons. (A) Expression of ELAV/Hu RBP members across a timecourse of directed differentiation of human iPSCs into neurons [67]. ELAVL1 (HuR) is ubiquitous and is downregulated upon neuronal specification, while ELAVL2-4 (HuB-D) are upregulated upon neuronal commitment. (B) K-means clustering identifies a cohort of genes with monotonically increasing *Ψ* across the neuronal differentiation timecourse, i.e. genes with neural-restricted (NR) alternative processing. (C-D) Collective behavior of NR-ALE (C) and NR-APA (D) genes across the neuronal differentiation timecourse. (E-F) Average behavior of NR-ALE (E) and NR-APA (F) genes across the neuronal differentiation timecourse. (G-H) Both NR-ALE (G) and NR-APA (H) genes specifically exhibit enrichment of uridine downstream of polyadenylation (pA) sites that are bypassed in neurons, compared to pA sites in other ALE and TUTR genes expressed in neurons. (I-J) The 100 nt regions downstream of NR-ALE (I) and NR-TUTR pA sites bypassed in neurons are highly enriched for ELAV/Hu RBP binding sites.

We followed this up using de novo motif searches. Interestingly, the top significant and most frequent (2/3 of loci) hit downstream of internal ALE termini that were bypassed in the presence of ELAV/Hu factors was a U-rich motif that closely resembled the consensus for ELAV/Hu factors (**Figure 5H**). Such a motif was either not found in other PAS categories (e.g., downstream of pA sites that were internal to 3’ UTRs of these genes, or downstream of the distal ALE termini of these genes), or was found at low frequency (9-12%) and with a different U-rich features (e.g., downstream of some classes of termini in genes unaffected by ELAV/Hu factors) (**Supplementary Figure 7**). We note that this motif is very similar to the motif recovered at cassette exons that appear to be directly regulated (excluded) by ELAV/Hu RBPs, from both GOF and LOF datasets (**Figure 2L-M**).

We performed similar analyses using larval CNS genes whose distal ALE usage was suppressed in *elav/fne* mutants compared to from *Canton-S* (wt). Here, we focused on genes expressed in CNS, which are only partially overlapping with S2-expressed genes, and thus reflect an independent analysis. The overall picture was quite similar. All CNS genes that are processed into multiple ALE isoforms exhibit somewhat weaker PAS at the termini of the internal ALE isoforms (**Supplementary Figure 6C**), but the major characteristic of ALE loci dependent on endogenous Elav/Fne is the presence of elevated uridine content downstream of pA sites (**Figure 5I-K**). Again, de novo motif searches show a high-frequency (44%) of U-rich motifs that closely resemble Elav binding sites downstream of internal ALE termini that are aberrantly expressed in *elav/fne* mutants (**Figure 5L**), whereas the frequency of such U-rich motifs downstream of other internal ALE termini is much lower (13%) (**Supplementary Figure 7**).

Taken together, these features extend the strategy of Elav-mediated pA bypass for distal ALE splice isoform switching, from *ewg* and *nrg* to many dozens of *Drosophila* loci. Moreover, we broaden this from Elav, thought to be the only nuclear ELAV/Hu family member in *Drosophila*, to other members. We show that in ectopic settings, all three members are able to drive distal ALE isoform usage by inducing downstream nascent transcription in the chromatin fraction, following pA site inhibition. Moreover, our comparison of single *elav* and double *elav/fne* mutants indicate that Fne plays a major role in controlling ALE splicing.

### Functionally overlapping requirements for Elav and Fne during neural differentiation

Elav is the most widely-used neuronal marker in *Drosophila*[60,61], yet there are relatively few phenotypic studies of its mutants, especially during its early lethal stages. Amongst the few embryonic defects attributed to *elav* mutants are aberrant commissural axon projections across the midline [62]. However, *elav* embryonic defects were reported to be the same upon concomitant mutation of cytoplasmic ELAV/Hu RBPs [54], suggesting that they have limited functional overlaps.

Our recent studies provide an alternative interpretation. Since the Elav paralogs Fne and Rbp9 exhibit limited protein accumulation during embryogenesis [63], it is conceivable that genetic interactions are not expected to be manifest until larval stages. Notably, while Elav is an acronym of “embryonic lethal, abnormal vision” [64], we recently clarified that upon dissection from the embryonic eggshell, *elav[5]* deletion mutants can survive as first instar (L1) larvae, as can *elav/fne* double null L1 larvae [44]. However, these single and double mutant larvae exhibit extreme locomotor defects, indicating that at least Elav is required for normal function of the nervous system (Supplementary Videos 1-3). Moreover, while these factors are normally segregated subcellularly, the L1-CNS exhibits substantial coexpression of Fne (but not Rbp9) with Elav, and Fne becomes substantially re-localized to the nuclei of *elav-*null neurons [44]. These considerations underlie the extensive functional overlap of Elav and Fne for 3’ UTR alternative polyadenylation [44], as well as for internal splicing and ALE splicing (this study) in the L1-CNS.

Inspection of our genomic data provided hints that Elav and Fne might co-regulate neural differentiation. For example, amongst L1-CNS genes with defective alternative splicing in elav/fne double mutants, the top enriched gene ontology (GO) terms for biological processes included synaptic transmission, axonogenesis, and synaptic growth (**Supplementary Figure 8**). It might be argued that some neural categories might come up when analyzing CNS data. However, when analyzing genes mis-spliced upon misexpression of Elav/Fne/Rbp9 in S2 cells, nervous system development still comes up as one of the top biological processes (**Supplementary Figure 8**). This suggests that ELAV/Hu RBPs are intimately linked to regulation of neural genes even when manipulated in a completely ectopic context. Several common categories of GO molecular functions are enriched amongst genes mis-spliced in elav/fne L1-CNS and upon gain-of-function of ELAV/Hu RBPs in S2 cells, including microtubule/tubulin binding proteins, actin binding factors, both of which include many factors involved in axonogenesis, as well as kinases and GTPases/GEFs (**Supplementary Figure 8**).

In light of these data, we analyzed whether *elav/fne* mutants exhibit phenotypic enhancement over *elav* or *fne* single mutants in the L1 CNS. We analyzed BP102, which labels CNS commissures and connectives [65], and was unaffected in *Δfne* (**Figure 6A-B**). Consistent with the *elav[5]* embryonic phenotype [62], these mutants exhibited midline crossing and occasional longitudinal breaks in the larval ventral nerve cord (**Figure 6C**). However, BP102 staining was much more severely disorganized in *elav/fne* double mutant CNS, with profound crossing, barely recognizable longitudinal organization, and general reduction of signal (**Figure 6D**). Mab 22C10 recognizes the Map1B homolog Futsch, which associates with microtubules and labels all axonal projections [66] (**Figure 6E**). *Δfne* mutants exhibit normal patterns (not shown), while *elav[5]* displays a mild reduction in 22C10 reactivity but the major nerves are present, if somewhat reduced (**Figure 6F**). However, 22C10 staining in *elav/fne* mutants was markedly compromised (**Figure 6G**). Overall, even though *elav/fne* double mutants resemble *elav* single mutant in embryos [54,62], these ELAV/Hu members exhibit strong phenotypic synergy in the larval CNS.

### Evidence that neural programs of mammalian distal ALE splicing and 3’ UTR lengthening are coordinated by ELAV/Hu RBPs

Our recognition that distinctive programs of neural ALE and APA are both mediated by ELAV/Hu factors in *Drosophila* inspired us to search for evidence of analogous regulatory programs during mammalian neural differentiation. To do so, we took advantage of recent directed human iPSC-neuron directed differentiation datasets [67]. These analyses comprise 9 timepoints including 3 stages of iPSC culture in “accelerated dorsal” media (days 2, 6 and 9), the neural precursor cell (NPC) stage (day 15), the neural rosette stage (21), and several timepoints following neural specification (days 49, 63 and 77), during which cells were co-cultured with rat astrocytes to facilitate terminal maturation.

Of the four mammalian ELAV/Hu members, ELAVL1/HuR is ubiquitously expressed, while ELAVL2-4/HuB-D are enriched in neurons. Accordingly, during iPSC->neuronal differentiation, *ELAVL1* transcripts are detected throughout, but are downregulated during the transition from neural rosettes to neurons. Concomitant with this, *ELAVL2/3/4* are all upregulated at this same transition, as neurons are first specified (**Figure 7A**).

To quantify APA across the neural differentiation time course, we used LABRAT (https://github.com/TaliaferroLab/LABRAT). LABRAT takes in RNAseq data and assigns psi (*Ψ*) values to genes with *Ψ* = 0 indicating exclusive usage of gene-proximal (upstream) polyadenylation sites and *Ψ* = 1 indicating exclusive usage of gene-distal (downstream) polyadenylation sites. We limited our analysis to genes that contained only two polyadenylation sites. With the neural differentiation time course, we used k-means clustering to identify genes with steadily increasing *Ψ* values throughout the differentiation. We termed these genes as having neural-restricted (NR) alternative processing. These include 706 genes with shifts towards longer 3’ UTRs within terminal 3’ UTRs (TUTR), 269 genes with shifts towards distal alternative last exons (ALE) (**Figure 7B)**. From these, we defined two substantial classes of NR genes, which directionally express more distal ALEs or lengthened TUTR isoforms (**Figure 7C-D**). Both classes of NR-ALE and NR-APA genes exhibited isoform shifts during the transition from neural rosettes into post-mitotic neurons, concordant with upregulation of *ELAVL2/3/4*. The remainder of genes that were expressed at the end of the timecourse (**Figure 7B**) were used as controls, which collectively exhibit unchanged or lower *Ψ* values in differentiated neurons compared to undifferentiated samples (**Figure 7E-F** and **Supplementary Figure 9**).

We analyzed the sequences downstream of NR-ALE and NR-TUTR proximal polyadenylation sites, and found that both exhibited significantly higher U-content than their respective controls (**Figure 7G-H**). This was likely attributable to binding by neuronal ELAV/Hu proteins, because de novo motif analysis revealed high frequency and highly significant occurrences of characteristic ELAV/Hu binding sites downstream of both NR-ALE and NR-TUTR proximal polyadenylation sites (**Figure 7I-J**).

Altogether, these bioinformatic analyses provide evidence for coordination of distal ALE switching and terminal 3’ UTR lengthening in mammalian neurons by neuronal ELAV/Hu RBPs, indicating that these two broad strategies of mRNA processing control in neurons are conserved across metazoans.

## Discussion

### Combined activities of *Drosophila* ELAV/Hu RBPs coordinately specify multiple aspects of mRNA processing

Programs of alternative splicing and alternative polyadenylation are orchestrated by multiple RBPs [1,2]. Several RBPs were reported to impact both types of mRNA processing events, but in most cases there does not appear to be substantial coordination in the directionality of alternate isoform generation. That is, upon manipulation of RBPs such as MBNL [29,68] or Nova [22,69,70], one can document both exon inclusion and exclusion events, as well as 3’ UTR lengthening and shortening events, but these seem not to be directionally correlated. Nevertheless, distal ALE and downstream tandem APA usage were reported to be correlated in mammals [26], with directionality toward more distal/longer isoforms in neurons. However, the underlying mechanisms were not specifically defined.

Regulation of distal ALE switching in the *erect wing* (*ewg*) gene was shown to be regulated by *Drosophila* Elav, which binds an extended region bearing bearing multiple U-rich motifs distal of the proximal cleavage site to inhibit proximal ALE 3’-end processing [40,71]. Likewise, regulation of APA was shown for all four Hu proteins in suppressing an intronic polyA site in the calcitonin/CGRP gene and HuR autoregulates by APA [39,72,73]. In addition, HuR regulates 3’-end processing of several membrane proteins [74]. Given the predominant neuronal expression of many ELAV/Hu members, these proteins are candidate regulators of CNS-specific 3’ UTR extensions. Elav was reported to mediate neural 3’ UTR extensions of certain genes [38], but the breadth of Elav involvement in the neuronal APA landscape was not determined until recently.

In concurrent work [44], we established that the three *Drosophila* ELAV/Hu members (Elav/Fne/Rbp9) are individually sufficient to induce the neural extended 3’ UTR landscape, and that endogenous overlapping activities of *Drosophila* Elav and its paralog Fne are critical to determine the extended 3’ UTR landscape of the larval CNS. Here, we extend this by showing that all three ELAV/Hu members are also able to specify hundreds of alternative splicing events, including alternative cassette exons and alternative last exons. Once again, we show the *in vivo* relevance of this, by demonstrating that dual deletion of *elav* and *fne* causes reciprocal changes to splice isoform accumulation. Notably, we are able to unify the rationale for three neuronal mRNA processing programs, in that the usage of isoforms associated with ELAV/Hu binding is suppressed. First, exons that are skipped in the presence of ELAV/Hu RBPs (either in S2 cells ectopically expressing Elav/Fne/Rbp9 compared to control S2 cells, or in wildtype L1-CNS compared to *elav/fne* double mutant L1-CNS) are enriched for flanking ELAV/Hu binding sites, but exons that are preferentially included do not. These sites are preferentially located upstream of skipped exons, although they are also enriched downstream of these regulated exons. Second, switching to distal alternative last exons is globally regulated by ELAV/Hu factors, in both GOF and LOF settings. Here, the relevant binding sites are preferentially located downstream of suppressed, proximal, cleavage sites, thereby permitting downstream transcription and splicing into distal ALEs. Third, lengthening of terminal 3’ UTRs is also globally regulated by ELAV/Hu factors, again in both GOF and LOF settings. Here, the logic is analogous to that of distal ALE switching, with binding downstream of suppressed pA sites.

Although mammalian ELAV/Hu factors were originally considered as post-transcriptional and translational regulators [35,75,76], select cases of their roles in mRNA processing were later recognized, including in splicing of specific targets [49,77] and APA regulation of ELAVL1/HuR by its neuronally restricted paralogs ELAVL2/3/4 (HuB/C/D) [72,73]. More recent genomic analyses of mammalian ELAV/Hu factors, including ELAVL1/HuR [45,46] and ELAVL2/3/4, now support the notion that these factors collectively play broad roles in alternative splicing, including both exon inclusion and exclusion [19,47]. However, relatively little has been examined with respect to ALE regulation or APA regulation. As all ELAV/Hu factors are likely to shuttle [78], it is worth bearing in mind that multiple ELAV/Hu factors may play roles in mRNA processing in mammals. Our study reveals that mammalian distal ALE switching and terminal 3’ UTR lengthening are correlated programs during neuronal differentiation, and are associated with similar layouts for enrichment for ELAV/Hu binding sites downstream of suppressed pA sites. These data provide a framework for analyzing combinatorial and correlated control of the mammalian neural transcriptome by ELAV/Hu RBPs.

### Biology of neuronal Elav/Hu factors

Certain members of the ELAV/Hu family are broadly expressed (e.g. vertebrate ELAVL1/HuR), and we recognized that seemingly neural-specific *Drosophila* Elav is actually ubiquitously expressed at a low level but post-transcriptionally repressed outside of the nervous system [79]. Nevertheless, a defining feature of this family is that most ELAV/Hu members in *Drosophila* and vertebrates are strongly elevated or restricted to neurons [80]. All three *Drosophila* members (Elav, Fne, Rbp9) are highly enriched in neurons, as are three of the four vertebrate family members (ELAVL2/3/4). Consistent with these expression patterns, ELAV/Hu members have been documented to have diverse neural phenotypes in different organisms. For example, mutants in *Drosophila elav* are lethal, and hypomorphic and/or conditional mutants exhibit severe defects in eye morphogenesis, neural differentiation, and locomotion, amongst other neural-related phenotypes. The other *Drosophila* family members are viable, but loss of *rbp9* compromises the blood-brain barrier [81] and deletion of *fne* causes male courtship defects [82]. In mice, knockout of *HuD* causes behavioral defects [83] while *HuB/D* double mutants exhibit seizures [19]. Zebrafish knockouts of HuD exhibit neural differentiation and behavioral defects [84], and a *C. elegans* Elav homolog is required for synaptic physiology [85]. Overall, ELAV/Hu proteins are implicated in a variety of neural gene regulation and behavioral regimes [86,87].

Our findings of substantial functional overlap by *Drosophila* ELAV/Hu factors is perhaps surprising, because they are normally located in distinct subcellular compartments and were not thought to exhibit mutant genetic interactions with respect to the embryonic nervous system [54]. However, it has been shown that the three ELAV/Hu paralogs have certain overlapping genetic activities [54], and our recent work [44] and current analysis affirm that they have substantially overlapping regulatory effects in ectopic settings. This is indeed relevant to *in vivo* neuronal biology, since while *fne* mutants have barely any phenotype on their own, we show that they strongly enhance the molecular and neuronal differentiation defects of *elav* mutant CNS. We can resolve the prior negative genetic interaction data [54] with these data by considering two factors. First, onset of Fne protein accumulation does not begin until late embryogenesis, while Elav protein appears much earlier and accumulates to much higher levels in neurons. Second, not only is Fne normally cytoplasmic, Elav actively excludes nuclear Fne by antagonizing inclusion of an *fne* microexon that directs a nuclear isoform [44]. Together, these data support the notion that Elav and Fne direct overlapping programs of neural mRNA processing that drive neural differentiation. We show that neuronal differentiation in double mutant L1-CNS is highly disrupted, much more so than either single mutant (although *elav* mutants clearly exhibit previously documented defects). Since we show that ectopic Rbp9 is capable of inducing similar cassette exon splicing, distal ALE switching, and APA lengthening programs as Elav and Fne, the collected data suggest that it may be productive to examine further double or triple mutant interactions amongst ELAV/Hu family members. This may require creative conditional genetics to achieve the requisite conditions, especially in pupal and/or adult stages, when Rbp9 is expressed at much higher levels in the nervous system [44]. We further suggest that concurrent knockout of the three neuronal ELAV/Hu members in mammals, perhaps in conjunction with loss of ubiquitous ELAVL1/HuR, will be critical to fully reveal their combined impacts on mammalian splicing and APA.

## Materials and Methods

### Identification and quantitation of alternative splicing events from RNA-seq

Total RNA-seq libraries were prepared from S2 cells transfected with actin-based ELAV/Hu family constructs (Flag-HA tagged-Elav/Fne/Rbp9) or mutant counterparts bearing point mutations in all three RRM domains of each factor (3xMut). We used the TruSeq Stranded Total RNA Library Preparation Kit (Illumina) with 2 μg input RNA as starting materials. Final cDNA libraries were sequenced on Illumina HiSeq-1000 sequencer with PE-100 mode. RNA-seq libraries of wildtype, *elav[5]* and *elav[5]/fne[Δ]* mutant L1-CNS were previously described [44].

To identify and quantify alternative splicing events, we used rMATS (v 3.2.5) [52] with - novelSS 1 parameter to detect unannotated splice sites. FDR<0.05 and ΔPSI>0.3 junction-spanning read counts for each alternative splicing events, including alternative 5′ splice site (A5SS), skipped exon (SE), mutually exclusive exons (MXE), retained intron (RI), and alternative 3’ splice site (A3SS) were used for downstream analysis.

### Identification and quantitation of alternative last exon events from 3’-end sequencing

3’-end sequencing libraries of both S2 and L1-CNS samples were previously described [44]. We mapped the sequenced reads to the *Drosophila melanogaster dm6* reference assembly and resulting reads were clustered and quantified within a 25 bp window as described [59].

To quantify alternative last exon usage by 3’-end cluster reads, we first identified genomic coordination of a specific 3’ UTRs generated by alternative last exon events (Unique 3’UTR). The unique 3’ UTRs which overlap the intronic regions of another isoform were identified, and the longest 3’ UTR was selected among the isoforms with the same 3’UTR start sites. Sum of the 3’-end cluster counts from the longest unique 3’ UTR was used for ALE events quantification.

To calculate ALE usage ratio, we first selected the unique 3’UTR that is used dominantly in the control, and calculated the utilization rates of the proximal unique 3’UTR and distal unique 3’UTR in both control and sample based on the dominant universal unique 3’UTR. ALE usage represents the value obtained by dividing the sum of universal unique 3’UTR and distal unique 3’UTR by the sum of universal unique 3’UTR and proximal unique 3’UTR, and the ALE usage ratio is the divided value of the ALE usage of sample and control.

### Relative strengths of internal ALE terminal ends

We calculate the bypass score for each ALE 3’-ends (internal ALE terminal ends) by percentage of sum of 3’-seq reads on the unique 3’UTR downstream of a given 3’end. The bypass ratio was calculated by comparing the bypass score of each ALE 3’-end between two samples in order to compare relative strength of ALE 3’-end. Bypass ratio 1 means that the ALE 3’-end has as much downstream last exons compared to the control and a ratio 0 means that the total amount of downstream last exon is not changed in both samples. Conversely, the unbypassed ratio is a measure of how much ALE 3’-end of a sample is not bypassed compared to the control.

### Motif analysis

We used MEME-suite (v 5.0.2) to discover *de novo* motifs arounds 50-nucleotide windows downstream of the cleavage site and 100-nucleotide windows of each up and downstream of the splicing site with default parameters [88]. Local enrichment of position weight matrix (PWM) of given motifs was analyzed by seqPattern R package in Bioconductor (https://bioconductor.org/packages/seqPattern/).

### *Drosophila* cell lines and transfections

*Drosophila melanogaster* S2R+ cells were maintained in Schneider’s Insect Medium supplemented with 10% heat inactivated FBS (VWR) and 1% Penicillin-Streptomycin (Thermo Fisher Scientific) at 25 °C. Cells were regularly passaged with the density of 2×10^6^/mL. Cell transfection was performed using Effectene (Qiagen) according to the manufacturer’s instructions. Transfections were done using cells at <20 passages using published expression constructs for Flag/HA-tagged constructs of wildtype and 3xRRM-Mut versions of Elav, Rbp9 and Fne, and Sxl, all in pAc5.1C vector [44].

### RNA extraction, reverse transcription PCR and real-time PCR

To process tissues, ∼30-40 whole 1st instar larvae or pools of dissected CNS were homogenized in 200 μL TRIzol using a Dounce tissue homogenizer (Thermo Fisher Scientific). L1 CNS dissections were performed in batches, with temporary storage on ice for <30 min before addition of TRIzol and storage at −80°C. TRIzol was used to extract RNA from S2R+ cells after pelleting washed cells.

RNA samples were treated with TurboDNase (Invitrogen) prior to reverse transcription. Poly(A)+ RNAs were enriched using Oligo(dT)25 magnetic beads (NEB). For reverse transcription using RNAs from S2R+ cells, 1 μg total RNA was used as input with a two-step reverse transcription using SuperScript III reverse transcriptase (Invitrogen) with either oligo(dT)18 priming or random priming, then 1/20 of RT product was used in a single PCR or real-time PCR reaction. For reverse transcription using RNAs from whole 1st instar larvae, 500 ng total RNA was used as input with a two-step reverse transcription with oligo(dT)18 priming, then 1/20 of RT product was used in a single PCR or real-time PCR reaction. To quantify ALE genes using total RNA from S2R+ cells, raw Ct values were normalized to *rpl32*. To quantify ALE genes using chromatin-associated RNA from S2R+ cells, raw Ct values were normalized to *roX2*. When quantifying ALE genes using 4sU-labeled RNA extracted from S2R+ cells, raw Ct values were normalized to *roX2*. When quantifying ALE genes using total RNA from whole 1st instar larvae, raw Ct values were normalized to *rpl14*. Primers used for RT-PCR and RT-qPCR analysis are listed in **Table S1**.

### Cell fractionation and isolation of nascent transcripts

72 hrs post-transfection, S2R+ cells were harvested and washed three times with PBS. Cell fractionation was performed as described [89]. Briefly, cells were lysed in hypotonic buffer (15 mM HEPES pH 7.6, 10 mM KCl, 5 mM MgOAC, 3 mM CaCl_2_, 300 mM sucrose, 0.1% Triton X-100, 1 mM DTT, 1X complete protease inhibitors) to rupture cell membranes and release nuclei. Nuclei were purified by centrifugation through sucrose cushion to remove intact cells, cell debris, etc., then further lysed with nuclear lysis buffer (10mM HEPES-KOH, pH 7.6, 100 mM KCl, 0.1 mM EDTA, 10% glycerol, 0.15 mM Spermine, 0.5 mM Spermidine, 0.1 M NaF, 0.1 M Na_3_VO_4_, 0.1 mM ZnCl_2_, 1 mM DTT, 1X complete protease inhibitors, 1 U/μL SuperaseIn). Then, 2X NUN buffer (25 mM HEPES-KOH pH 7.6, 300 mM NaCl, 1 M Urea, 1% NP-40, 1 mM DTT, 1X complete protease inhibitors, 1 U/μL SuperaseIn) was added to the suspension with 1:1 ratio to nuclear lysis buffer. After centrifugation, the supernatant comprises nuclear lysate while the pellet contains DNA/histones/Pol II-RNA containing nascent RNA transcripts. We saved 5% of each fraction for western blot analysis, and subjected the remainder to RNA extraction using TRIzol (Invitrogen).

### 4sU-labeling and 4sU-containing transcripts isolation

Cells were cultured in medium supplemented with 100 μM of 4sU for 1 hr before harvest. Total RNA was extracted using TRIzol. 4sU-labeled and pre-existing RNA populations were fractionated as described [90]. Briefly, 100 μg of total RNA was diluted in 1X Biotinylation buffer (100 mM Tris pH 7.4, 10 mM EDTA) with biotin-HPDP (1 μg/μl in DMF), incubated at room temperature on a rotator for 1.5 hrs. RNA was then extracted with Phenol:Chloroform:Isoamyl Alcohol and precipitated in EtOH for at least 2 hrs. RNA pellet was dissolved in 50 μl of nuclease-free H2O and denatured by incubation at 70 °C for 2 min. After chilling on ice, RNA was mixed with 50 μl of pre-washed Streptavidin C1 Dynabeads in 2X bind and wash buffer (10 mM Tris-HCl pH 7.5, 1 mM EDTA, 2 M NaCl, 1 U/μl SuperaseIn). The mixture was incubated on a rotator at room temperature for 1 hr. After incubation, beads were collected on a magnetic stand, and supernatant containing the pre-existing RNAs was discarded. The beads were washed three times with 0.5 ml 1x bind and wash buffer (5 mM Tris-HCl pH 7.5, 0.5 mM EDTA, 1 M NaCl) at 65 °C, followed by three washes with 1X bind and wash buffer at room temperature. After complete removal of the bind and wash buffer, beads were resuspended in 200 μl of 1x bind and wash buffer containing 100 mM DTT, and incubated at room temperature for 3 min to elute 4sU-labeled RNA. The elution process was repeated and the eluted RNA was precipitated as described above. The isolated RNA was used for RT-qPCR.

### *Drosophila* immunostaining

We used *Canton-S* as a wildtype reference, and the deletion alleles *elav[5]* and *Δfne*. The single *elav* mutant and *elav*/*fne* double mutants were maintained over *FM7, Dfd-GFP*. Wildtype and mutant CNS were dissected from 1st instar larvae in cold PBS using Dumont #5 forceps (Fine Science Tools). The CNS were fixed in 4% paraformaldehyde in PBS (Sigma Aldrich) at room temperature for 1 hr, followed by two 30’ washes in PBS with 0.2% Tween 20 (PBST). The following primary antibodies were used: rat anti-Elav (1:1000; DSHB), mouse anti-22C10 (1:100, DSHB), mouse anti-BP102 (1:100, DSHB). Samples were incubated with the primary antibody overnight at 4°C, followed by two 30’ washes in PBST. Secondary antibodies were goat anti-mouse IgG Alexa Fluor 546, and goat anti-rat lgG Alexa Fluor 633 (each 1:1000, Invitrogen). Samples were incubated in secondary antibodies for 1 day at 4°C with gentle rotation, followed by two washes in PBST, each 30 min. Samples were incubated with DAPI in PBS at room temperature for 1 hr before mounting. Imaging was performed on a Leica SP5 spectral confocal microscope using HCX PL APO 63X∼/0.70 and HCX PL APO 100X∼/1.25-0.75 lenses and processed using FIJI.

### Analysis of human differentiation time course samples

RNAseq data for human iPSC to mature neurons was downloaded from https://www.ncbi.nlm.nih.gov/bioproject/?term=PRJNA596331) [67]. Each sample is derived from one of several different donors and differentiated through multiple timepoints. To quantify 3’ end usage across neuronal differentiation, the LABRAT software package was used.

LABRAT is freely available for download here: https://github.com/TaliaferroLab/LABRAT/. LABRAT quantifies APA events by comparing the expression of the final two exons of every expressed transcript and calculating a *Ψ* value that relates relative APA site usage. To simplify LABRAT output, only gene models with two APA isoforms were considered. For these genes, *Ψ* values of 0 indicate exclusive usage of the proximal APA site while *Ψ* values of 1 indicate exclusive usage of the downstream APA site.

To define tandem UTR and ALE gene models, LABRAT observes the isoform structures at the 3’ end of a gene. If all APA sites are contained within the same exon, then the structure in tandem UTR. If all APA sites are contained within different exons, then the structure is ALE. If a gene has more than two APA sites, it is possible for the gene to fit into neither classification. In these cases, LABRAT assigns the gene to have a “mixed” structure. These genes were not considered in following analyses.

To define genes that change their APA usage through neuronal differentiation by using more downstream APA sites, we utilized standard k-means clustering. Genes with changing values across timepoints were clustered by their mean *Ψ* value revealing a distinct cluster of genes with increasing *Ψ*. These genes were termed as having “neuronally restricted” (NR) long isoforms. These include 706 genes with shifts towards longer 3’ UTRs within terminal 3’ UTRs (TUTR), 269 genes with shifts towards distal alternative last exons (ALE), and 15 genes with mixed TUTR/ALE profiles; the small latter class was not considered further in our analysis. A control group of genes was defined as being expressed in the later differentiation time points (Day 47, 63, and 77) but either had no change in *Ψ* or showed decreasing *Ψ* values across the differentiation time course.

Sequences flanking the proximal PAS (100 nucleotides upstream and downstream) were analyzed for each NR and control gene. Uridine content was plotted and compared using Wilcoxon rank-sum tests and considered significant with a p < 0.05. Additionally, these sequences were analyzed for ELAVL PWM motif matches of greater than 80%. ELAVL motifs were obtained from cisbpRNA database [55] and RNA bind-n-seq results [91]. Multiple motifs represent each of the 4 protein family members and motifs can be quite similar. Motifs with sufficient match to a sequence were centered onto a single nucleotide. If a similar motif matched at the exact location, it was not counted twice. Every motif count for an ELAVL protein was summed across nucleotide positions for the NR and control gene groups. Motif frequency within the groups was plotted and compared using Wilcoxon rank-sum tests and considered significant with a p < 0.05.

## Acknowledgments

Work in E.C.L.’s group was supported by the National Institutes of Health (R01-NS083833 and R01-GM083300) and MSK Core Grant P30-CA008748. Work in J.M.T’s group was supported by the National Institutes of Health (R35-GM133885), a Predoctoral Training Grant in Molecular Biology (RG) (T32-GM008730), and by the RNA Bioscience Initiative at the University of Colorado Anschutz Medical Campus.

## Supplementary Figure Legends and Tables

**Supplementary Figure 1**. Analysis of changes in alternative splicing events following manipulation of *Drosophila* ELAV/Hu RBPs. (A) Gain-of-function of ELAV/Hu RBPs in S2 cells. The effects of each WT ELAV/Hu RBP (Elav/Fne/Rbp9) were compared to their RNA binding defective counterparts (3X-Mut). The (B) Loss of function of ELV/Hu RBPs in 1st instar larval CNS (L1-CNS). Pairwise comparisons between wildtype *Canton-S*, single *elav[5]* null and double *elav[5]/Δfne* null mutants. The stacked bar plots on the left categorize the numbers of all differential alternative splicing events involving internal exons. The middle plots and right volcano plots focus only cassette exons, which comprise the most numerous category of alternative splicing event affected by ELAV/Hu RBP manipulation.

**Supplementary Figure 2**. Examples of exon skipping and exon inclusion promoted by *Drosophila* ELAV/Hu RBPs. (Top) Three examples of genetic sufficiency and necessity for exon skipping promoted by ELAV/Hu RBPs. GOF of wildtype Elav/Fne/Rbp9 (but not their RNA-binding defective counterparts) promotes skipping of an alternative cassette exon in S2 cells, while combined LOF of *elav/fne* results in inclusion of the same exon in L1-CNS. (Bottom) Three examples of genetic sufficiency and necessity for exon inclusion promoted by ELAV/Hu RBPs. GOF of wildtype Elav/Fne/Rbp9 (but not their RNA-binding defective counterparts) promotes inclusion of an alternative cassette exon in S2 cells, while combined LOF of *elav/fne* results in exclusion of the same exon in L1-CNS.

**Supplementary Figure 3**. Flanking intronic features associated with exon exclusion and skipping regulated by *Drosophila* ELAV/Hu RBPs. Shown are nucleotide frequencies and de novo motif searches from flanking intronic regions of regulated cassette exons from ELAV/Hu RBP GOF in S2 cells (top) or *elav/fne* LOF in L1-CNS (bottom). These are separated according to exons that are either included or excluded in the corresponding manipulated condition, bearing in mind that exons that are included in elav/fne LOF correspond to ones that are excluded in the wildtype condition (since these factors are normally expressed in CNS). We observe overt enrichment of upstream intronic uridine (blue oval) amongst introns that are excluded by misexpression of Elav/Fne/Rbp9 (EFR) in S2 cells or are included in elav or elav/fne knockout L1-CNS. This is associated with strong enrichment of ELAV/Hu-type binding sites from de novo motif searches in their upstream flanking introns, as well as a lesser but still significant enrichment of such sites in flanking downstream intronic regions. Exons that are included by ELAV/Hu RBP activity do not show such flanking uridine enrichment or ELAV/Hu-type binding sites, but instead show enrichment of downstream flanking adenosine (red oval).

**Supplementary Figure 4**. Additional examples of distal alternative last exon splicing usage promoted by *Drosophila* ELAV/Hu RBPs. These four genes generally share the features of the distal ALE isoforms that are (1) preferentially expressed in head compared to other tissues (detection in carcass may correspond to presence of ventral nerve cord in these dissected samples), (2) are developmentally induced during the timecourse of embryogenesis, (3) are induced in S2 cells upon transfection of wildtype Elav/Rbp9/Fne but not their RRM-mutant (Mut) counterparts, and (4) are expressed in dissected wildtype *Canton-S* and *elav[5]* null L1-CNS, but not in *elav[5]/Δfne* double mutant L1-CNS. Tracks are labeled as to 3’-seq or RNA-seq data.

**Supplementary Figure 5**. Validation of terminal 3’ UTR extensions respond to ELAV/Hu RBPs. S2 cells were transfected with wildtype (WT) or mutant (Mut) versions of Elav/Rbp9/Fne; the latter constructs contain point mutations in all three RRM domains. qPCR of total RNA samples for universal (uni) and extension (ext) 3’ UTR amplicons shows that the 3’ UTRs of *tai* and *ctp* are specifically extended by ectopic wt ELAV/Hu RBPs. Measurements were normalized to *rpl32*.

**Supplementary Figure 6**. Analysis of polyadenylation signals (PAS) within ALE genes. (A) Schematic of ALE gene models and designation of PAS locations. (B) PAS types in S2-expressed genes with a single 3’ end. A strong majority of genes utilize the canonical AAUAAA PAS or one of its top two *Drosophila* variants (AUUAAA/AAUAUA). (C) In S2 cells, ALE-expressed genes exhibit lower frequency of canonical PAS at their proximal ALE 3’ ends than the single-end gene reference. However, these are not substantially different between loci that are bypassed in the presence of ectopic ELAV/Hu RBPs and ones that are not. (D) In L1-CNS, ALE-expressed genes exhibit lower frequency of canonical PAS at their proximal ALE 3’ ends than the single-end gene reference. However, these are not substantially different between loci that are bypassed in the presence of ectopic ELAV/Hu RBPs and ones that are not. Note that in both cell/tissue cohorts, “other PAS” exhibit overall lower quality features, which is consistent with these collectively including a portion of biochemically valid although potentially less biologically critical PAS.

**Supplementary Figure 7**. ELAV/Hu-type binding sites are highly enriched downstream of pA sites of proximal ALE models that are bypassed by ELAV/Hu RBPs. Shown are results of de novo motif analysis from 50 nt regions downstream of various cohorts of polyadenylation (pA) sites from ALE gene models of loci expressed in S2 cells or L1-CNS. Significant motifs recovered at >10% frequency are shown. (Top) In S2 cells, proximal ALEs that were prone to being switched to distal ALEs in the presence of wt Elav/Fne/Rbp9 are highly enriched for downstream ELAV/Hu-type binding sites. Note that there is also enrichment for similar sites downstream of unchanged proximal ALE pA sites, albeit at a much lower frequency, potentially indicating that ELAV/Hu RBPs may be involved in their regulation but were insufficient to mediate their switching in these experimental tests. (Bottom) In L1-CNS, the only motif recovered downstream of proximal ALE PAS that became aberrantly switched from distal ALE usage in elav/fne double mutants was the ELAV/Hu-type site. Similar to the S2 tests, we also observed significant, but much less frequent, enrichment of similar sites downstream of proximal ALE PAS that were not affected in elav/fne mutants. Again, this may indicate that the involvement of ELAV/Hu RBPs is broader than functionally detected in these data, either because of the need for the triple mutant or because of other parallel regulatory mechanisms.

**Supplementary Figure 8**. Gene ontology (GO) analysis of genes with ELAV/Hu RBP-regulated alternative mRNA processing. The top row shows genes whose processing is altered by ELAV/Hu gain-of-function in S2 cells, by transfection of Elav/Fne/Rbp9-wt compared to their RNA binding defective counterparts. The bottom row shows genes whose processing is altered by ELAV/Hu loss-of-function in dissected L1-CNS, by comparison of control Canton-S to single *elav* mutant or *elav/fne* double mutant. GO analysis for three groups of genes are shown, switches of distal alternative last exons (ALE), alternative splicing (AS) of internal cassette exons, and lengthening of terminal 3’ UTRs by alternative polyadenylation (APA). Because the numbers of ALE targets is the smallest, it generates the least significant GO outputs. We highlighted a number of AS and APA GO terms that were overlapping between the S2 and L1-CNS datasets, which included several similar biological process terms and molecular function terms. Note that even when manipulating ELAV/Hu RBP factors in S2 cells, which are of macrophage origin, the regulated genes define many GO terms relevant to neuronal development or differentiation.

**Supplementary Figure 9**. Psi value changes for genes with alternative 3’ ends across human iPSC-neuronal directed differentiation. Each row of a timecourse plot shows the distribution of LABRAT calculated psi values across loci relative to the starting day 02 dataset. The top plots of genes with neural-restricted (NR) distal ALE-switching (left) or terminal 3’ UTR (TUTR) lengthening (right) are the same as in Figure 7C-D and shown for reference. The bottom plots comprise the control sets of ALE and TUTR gene used for analysis in Figure 7.

